# RT-QuIC detection of CWD prion seeding activity in white-tailed deer muscle tissues

**DOI:** 10.1101/2021.03.22.436515

**Authors:** Manci Li, Marc D. Schwabenlander, Gage R. Rowden, Jeremy M. Schefers, Davis Seelig, Peter A. Larsen

**Affiliations:** Department of Veterinary and Biomedical Sciences, University of Minnesota, MN 55108; Minnesota Center for Prion Research and Outreach, College of Veterinary Medicine, University of Minnesota, MN 55108; Veterinary Population Medicine Department, Veterinary Diagnostic Laboratory, University of Minnesota, MN 55108; Department of Veterinary Clinical Sciences, University of Minnesota, MN 55108

**Keywords:** Chronic wasting disease, *Odocoileus virginianus*, real-time quaking-induced conversion, skeletal muscle, venison

## Abstract

Chronic wasting disease (CWD) is a prion disease circulating in wild and farmed cervid populations throughout North America (United States and Canada), Europe (Finland, Norway, Sweden), and South Korea. CWD is an immediate threat to cervid heritage and the disease is causing substantial economic losses across multiple sectors. In North America, hunting and farming industries focused on the processing and consumption of white-tailed deer (WTD) venison are particularly vulnerable to CWD, as millions of WTD are consumed annually. Real-time quaking-induced conversion (RT-QuIC) is a highly sensitive assay amplifying misfolded CWD prions *in vitro* and has facilitated CWD prion detection in a variety of tissues and excreta. To date, no study has comprehensively examined CWD prion content across bulk skeletal muscle tissues harvested from individual CWD infected WTD. Here, we use RT-QuIC to quantify prion-seeding activity in a variety of skeletal muscles from both wild and farmed CWD-positive WTD. We successfully detected CWD prions in muscles commonly used for consumption (e.g., backstrap, tenderloin, etc.) as well as within tongue and neck samples of WTD. Our results help to establish the utility of RT-QuIC for monitoring CWD prions in venison and suggest that the method is useful for preventing CWD prions from entering animal and human food chains. Moreover, our work indicates that CWD prions are more widely distributed across skeletal muscles of infected WTD than previously reported.

## Introduction

Chronic wasting disease (CWD) is an infectious and fatal prion disease transmitted among cervids, including white-tailed deer (WTD; *Odocoileus virginianus*), mule deer, elk, red deer, caribou, reindeer, and moose. The disease threatens all aspects of cervid heritage, and it is now prevalent in the USA, Canada, Korea, and Scandinavian regions [1]. A number of cervid-related multibillion-dollar economic sectors are negatively impacted by CWD, including both agricultural and hunting industries. As with other transmissible spongiform encephalopathies [2,3], CWD prion seeds (PrP^CWD^) consist of misfolded cellular prion protein (PrP^C^) which form β-sheet-rich amyloid fibrils through inducing conformational change and polymerization of native PrP^C^. The central nervous system (CNS) typically contains the highest load of prions in a terminally diseased animal in comparison to peripheral tissues and body excreta due to the abundance of PrP^C^ in nervous tissues [4].

Currently, CWD diagnosis relies on the identification of Proteinase K (PK)-resistant PrP^CWD^ by enzyme-linked immunosorbent assay (ELISA) and immunohistochemistry (IHC) [5]. In the past two decades, the detection of prion seeding-activity was greatly enhanced by highly sensitive methods involving amplification of protein misfolding *in vitro*, such as protein misfolding cyclic amplification (PMCA) and real-time quaking-induced conversion (RT-QuIC). PMCA uses rodent brain homogenates as the substrate to amplify misfolded prions and Western Blotting as the output [5–7]. RT-QuIC utilizes recombinant PrP^C^, commonly from rodent sources, as the substrate for prion amyloid formation, the real-time reporting of which is enabled by thioflavin T (ThT) binding and detection [8–10].

CWD prions have previously been identified in a variety of tissue types and excreta using RT-QuIC, such as the CNS, third eyelids, and feces [11–14]. Prior studies focused on the detection of CWD prions in skeletal muscle using immunodetection methods have produced mixed results [15–18]. PMCA was used to amplify CWD prions in hindlimb muscles from two WTDs [17]. Although RT-QuIC has the advantage over other methods for screening due to its high sensitivity and adaptability, no comprehensive reports are available for detecting CWD prions using RT-QuIC in skeletal muscle. Skeletal muscle tissues from CWD-infected deer contain infectious prions as determined in transgenic mice bioassay [18].

Recent studies have shown that there are compelling reasons to conclude that CWD poses a non-zero risk to a variety of mammals, including humans [1,19]. Challenge experiments using CWD-causing prions have shown that CWD can cause neurodegenerative disease in numerous species, including ferrets, mink, domestic cats, sheep, goats, cows, pigs, and squirrel monkeys [20]. *In vitro* experiments showed that CWD prions can convert human prion proteins into a misfolded and potentially disease-causing form [19]. For these reasons, as of 2020, both the FDA (Food and Drug Administration) and FSIS, USDA (Food Safety and Inspection Service, United States Department of Agriculture) consider venison from CWD-positive animals as adulterated and unsuitable for consumption [21,22]. However, at the time of this publication, there are no guidelines regarding venison-based detection of CWD and associated surveillance. This observation, combined with the limitations of existing CWD diagnostic tools (e.g., ELISA and IHC), has resulted in a situation whereby venison processing occurs without the knowledge of an animal’s CWD status, and it is estimated that at least 15,000 CWD positive cervids are consumed in the USA annually [1]. Underscoring this discussion is a well-documented 2005 exposure event in which over 200 participants at a Sportsmen’s feast consumed CWD-positive venison [23]. Current estimates indicate a 20-50% CWD prevalence rate in harvested white-tailed deer from focal areas of southern Wisconsin, however, only 1 out of 3 are tested for the disease [24]. Additionally, venison products have become a common ingredient in pet food (e.g. commercial cat and dog food). These observations indicate an emerging food-related risk to human and animal health. Therefore, the development of a scalable method for sensitive CWD-prion detection in venison is urgently needed. Research focused on the distribution of PrP^CWD^ in cervid skeletal muscle tissue is needed to develop a deeper understanding of potential risks.

Here, we examine the utility of RT-QuIC for the detection of CWD prions within a broad set of WTD skeletal muscle tissues, including those frequently used for both human and animal consumption. We report the RT-QuIC results for muscles sampled from the neck (*brachiocephalicus/sternocephalicus*) of wild WTD with known CWD status. Further, we investigated whether CWD prion deposition is limited to certain groups of muscles or if it is more generalized by using multiple WTD skeletal muscle groups across the body, including muscles from the tongue, forelimb (*suprascapularis*), backstrap (*longissimus dorsi*), tenderloin (*psoas major*), and hindlimb (*semimembranosus/semitendinosus*) from both wild and farmed CWD positive animals independently determined by ELISA and/or IHC.

## Results

### RT-QuIC detection of CWD prions in unilateral skeletal muscles from the neck of wild WTD

To test the PrP^CWD^ enrichment protocol detailed below (see methods), we processed unilateral muscles collected from the neck (*brachiocephalicus/sternocephalicus*) of 10 CWD-positive and 10 CWD-negative wild WTD (Table 1) and analyzed the resultant homogenates using RT-QuIC. We found that we could detect significant prion seeding activity in 8 out of 10 (80%) samples from 10 different CWD-positive animals (Table 1; Fig 1A) with relatively consistent fluorescent readings (Fig 1B). In contrast to animals with official CWD-positive test results, none of the muscle samples from CWD-not-detected animals showed statistically significant prion seeding activity in RT-QuIC (Fig 1B), despite one of eight wells crossing the threshold from a single animal (Fig 1D).

**Table 1.**
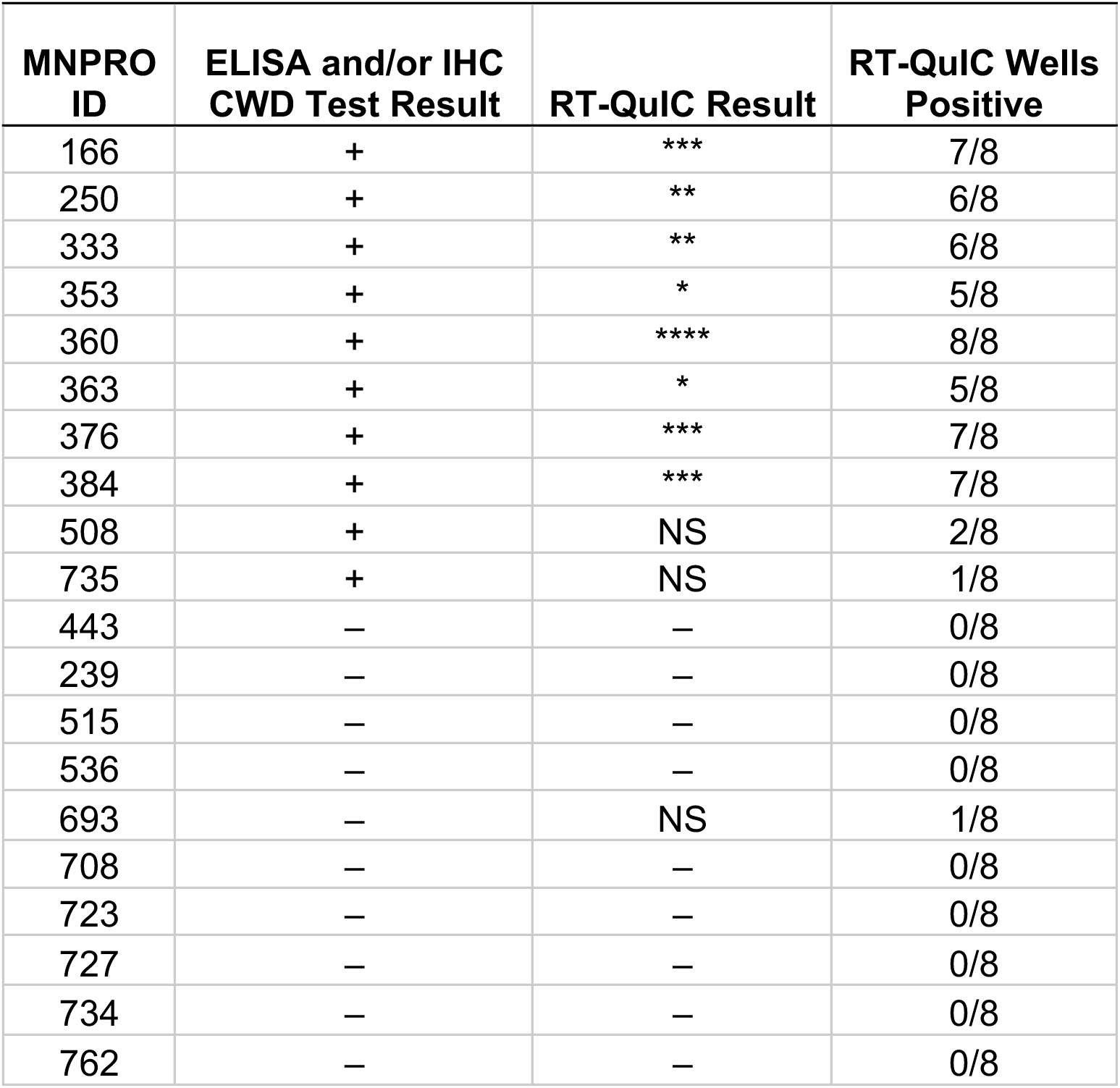
RT-QuIC results of WTD neck muscles. All animals were collected through Minnesota Department of Natural Resources agency culling operations and tested for CWD through official regulatory means (e.g., ELISA and/or IHC) based on retropharyngeal lymph node. NS, rate of amyloid formation is not 0 but not statistically significant from the corresponding negative controls; –, rate of amyloid formation is 0 in the given time period; ****, p < 0.0001; ***, p < 0.001; **, p < 0.01; *, p < 0.05. The freeze-thaw method was used for sample processing and RT-QuIC was performed at 45°C.

| MNPRO ID | ELISA and/or IHC CWD Test Result | RT-QuIC Result | RT-QuIC Wells Positive |
| --- | --- | --- | --- |
| 166 | + | *** | 7/8 |
| 250 | + | ** | 6/8 |
| 333 | + | ** | 6/8 |
| 353 | + | * | 5/8 |
| 360 | + | **** | 8/8 |
| 363 | + | * | 5/8 |
| 376 | + | *** | 7/8 |
| 384 | + | *** | 7/8 |
| 508 | + | NS | 2/8 |
| 735 | + | NS | 1/8 |
| 443 | – | – | 0/8 |
| 239 | – | – | 0/8 |
| 515 | – | – | 0/8 |
| 536 | – | – | 0/8 |
| 693 | – | NS | 1/8 |
| 708 | – | – | 0/8 |
| 723 | – | – | 0/8 |
| 727 | – | – | 0/8 |
| 734 | – | – | 0/8 |
| 762 | – | – | 0/8 |

**Figure 1.**
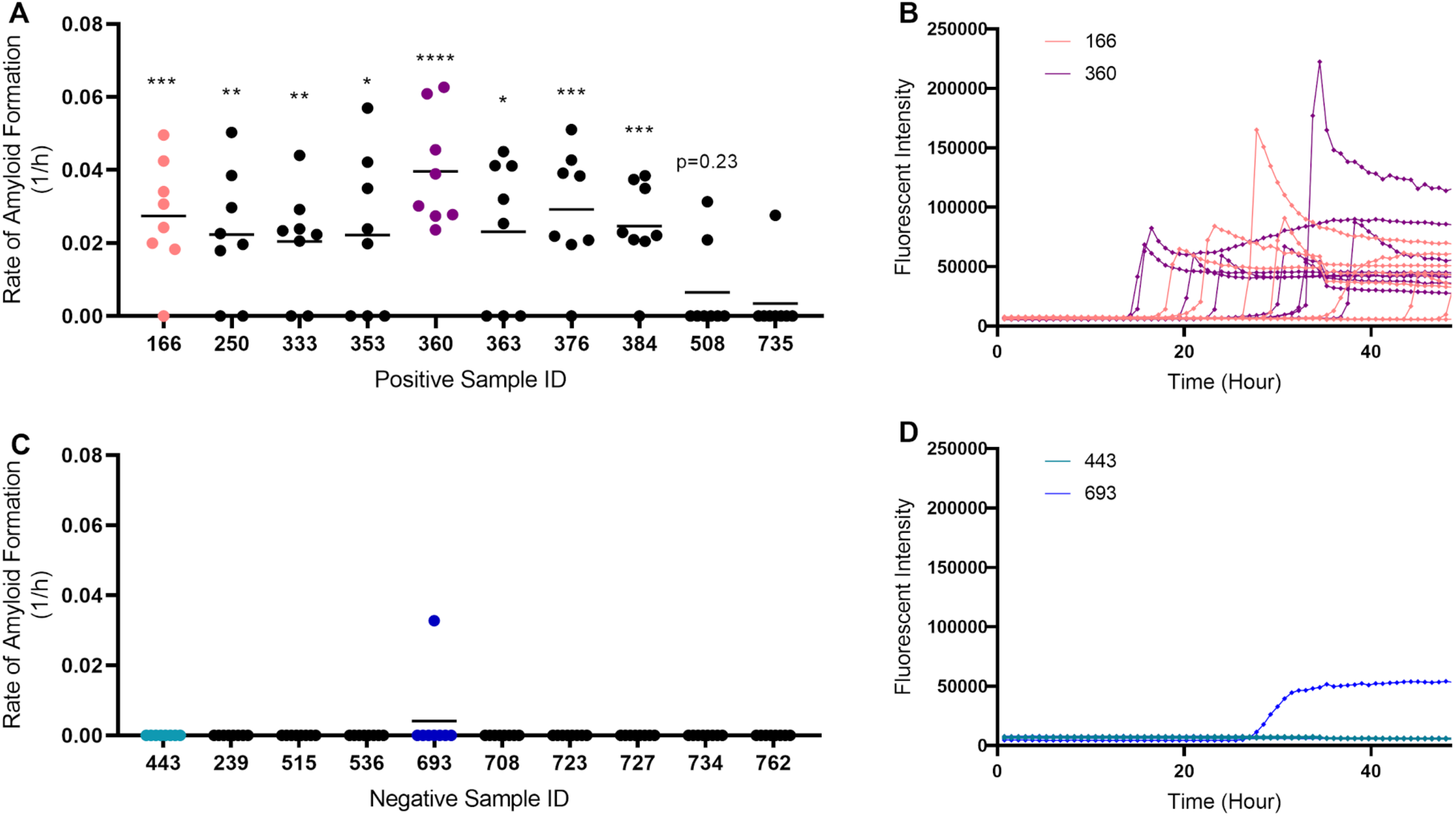
Detection of prion seeding activity in unilateral neck muscles from white-tailed deer. (A) Rate of amyloid formation (1/h) was plotted using data collected from CWD-positive animals. Statistical significance was obtained through comparing with rate of amyloid formation with the respective negative controls on the same plate (****,p < 0.0001; ***, p < 0.001; **, p < 0.01; *, p < 0.05). (B) Examples of real-time fluorescence readings from positive animals (sample numbers 166 and 360). (C) Rate of amyloid formation (1/h) from CWD-negative animals. (D) Examples of real-time fluorescence readings from negative animals (443 and 693); plotted as described in B and showing one of eight wells for sample 693 having amyloid seeding activity (not significant).

We then compared the rate of amyloid formation (RAF) among muscles, blood, and lymphoid tissues, all of which were processed using mechanical extraction methods; with methods and results of blood and lymphoid tissues reported by Schwabenlander et al. [25]. We note that although muscles appeared to have a lower RAF, animals could have a statistically positive RT-QuIC result for muscles and lymphoid tissues but not blood (e.g., animal 166; Fig 2A). In addition, the freeze-thaw method presented here routinely underestimates the RAF for consistent detection purposes. Using 10^−1^ dilution of the enriched homogenates, prion load would be considered as 100 to 1000 times lower than previous reports in muscles of BSE prion-inoculated hamster (Supplementary Fig 1) [26]. The difference is likely due to inhibitors of prion-seeding activity in muscle tissues rather than incomplete extraction. 10^−1^ dilution was chosen because of its consistency in producing results in different animals (Fig 2B). The optimal dilutions of each animal may differ, with dilution factors ranging from 0 to 2 (Fig 2B).

**Figure 2.**
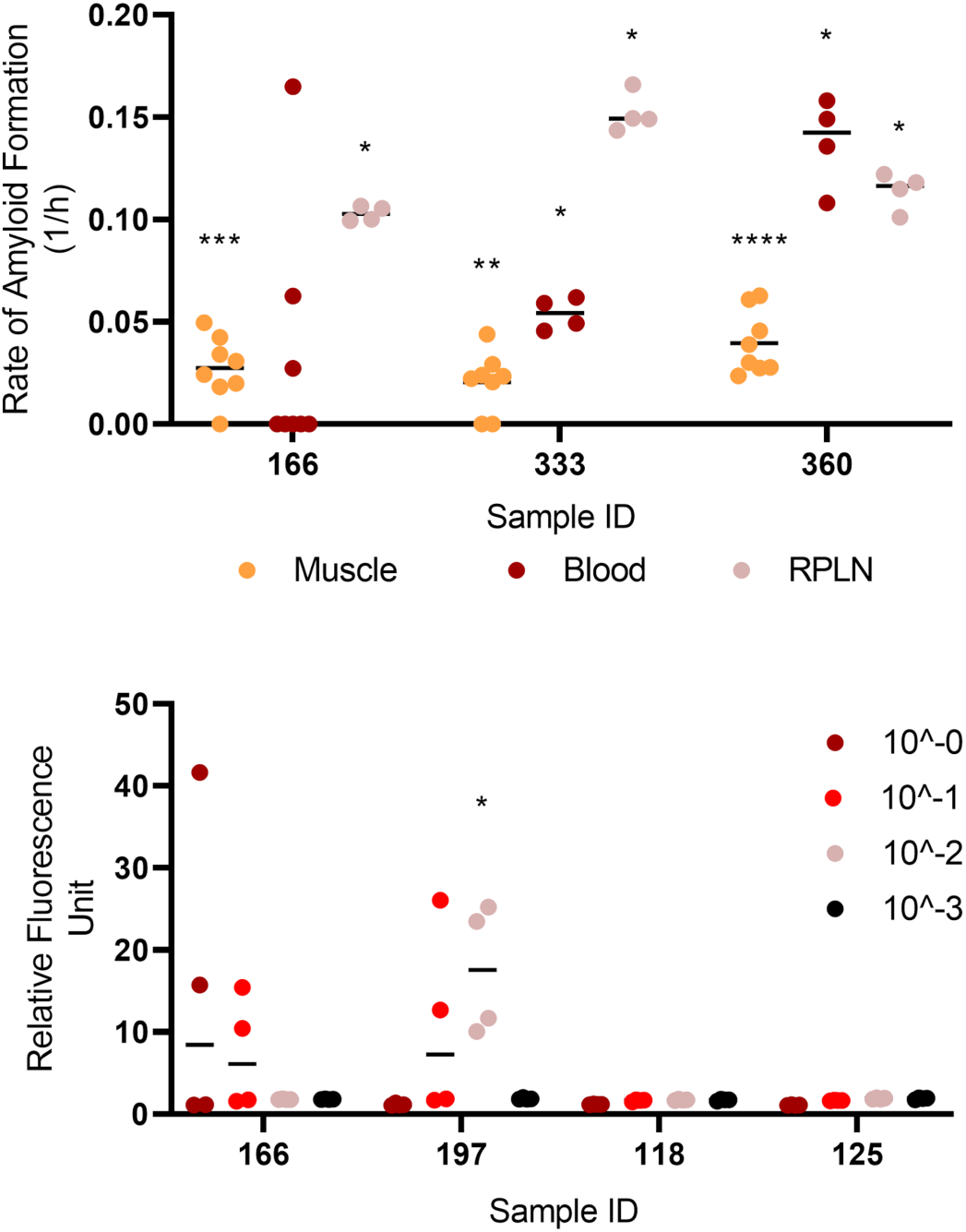
Comparison of prion-seeding activity in RT-QuIC. (A) Rate of amyloid formation was compared among neck muscle, blood, and lymphoid tissues of three CWD-positive animals. (B) Relative fluorescence units (RFUs) were compared among different dilutions of the enriched homogenates. RFUs were calculated using the last cycle of fluorescence reading divided by that of the initial reading. RFUs were compared between each sample and the negative control on the same plate. Statistical result for each sample compared with its respective negative control was indicated (****, p < 0.0001; ***, p < 0.001; **, p < 0.01; *, p < 0.05). RPLN, retropharyngeal lymph nodes.

### CWD prions found in muscles from the tongue, neck, mid-trunk, forelimb, and hindlimb of WTD

To investigate if the CWD prions are widely distributed in WTD skeletal muscles and whether the freeze-thaw method described above can be used to detect prions deposition in other skeletal muscles other than those from the neck, we used a set of muscle tissues from another 10 WTD, including the tongue, forelimb (*suprascapularis*), backstrap (*longissimus dorsi*), tenderloin (*psoas major*), and hindlimb (*semimembranosus/semitendinosus*). In the blinded run, we were able to detect at least one significantly RT-QuIC positive sample in all the muscle groups tested (Table 2; Fig 3). We conclude that PrP^CWD^ is more widely distributed throughout the skeletal muscles of WTD than previously reported (Fig 3B) [17].

**Table 2.** RT-QuIC results of various WTD muscle groups. All animals were tested for CWD through official regulatory means (e.g., ELISA and/or IHC) based on obex and/or retropharyngeal lymph node. NS, rate of amyloid formation is not 0 but not statistically significant from the corresponding negative controls; –, rate of amyloid formation is 0 in the given time period; ****, p < 0.0001; ***, p < 0.001; **, p < 0.01; *, p < 0.05. RT-QuIC analyses of forelimb (F), hindlimb (H), backstrap (B), tenderloin (Td), and tongue (Tg) muscles were performed with the researcher blinded to official CWD testing results (see methods). The freeze-thaw method was used for sample processing and RT-QuIC was performed at 45°C.

| ID | Region | MNPRO ID | ELISA and/or IHC CWD Test Result | RT-QulC Result | RT-QulC Wells Positive |
| --- | --- | --- | --- | --- | --- |
| 1 | F | 287 | + | - | 0/8 |
| 2 | H | 287 | + | NS | 3/8 |
| 3 | Tg | 287 | + | *** | 8/8 |
| 4 | B | 287 | + | - | 0/8 |
| 5 | F | 288 | + | NS | 1/8 |
| 6 | H | 288 | + | NS | 1/8 |
| 7 | B | 288 | + | NS | 1/8 |
| 8 | F | 289 | + | NS | 1/8 |
| 9 | H | 289 | + | - | 0/8 |
| 10 | B | 289 | + | - | 0/8 |
| 11 | F | 290 | + | - | 0/8 |
| 12 | H | 290 | + | NS | 2/8 |
| 13 | Tg | 290 | + | - | 0/8 |
| 14 | Td | 290 | + | - | 0/8 |
| 15 | H | 295 | + | NS | 1/8 |
| 16 | F | 295 | + | - | 0/8 |
| 17 | B | 295 | + | NS | 1/8 |
| 18 | H | 296 | + | - | 0/8 |
| 19 | F | 296 | + | - | 0/8 |
| 20 | B | 296 | + | - | 0/8 |
| 21 | H | 297 | + | NS | 1/8 |
| 22 | F | 297 | + | NS | 1/8 |
| 23 | B | 297 | + | - | 0/8 |
| 24 | H | 298 | + | NS | 1/8 |
| 25 | F | 298 | + | - | 0/8 |
| 26 | B | 298 | + | - | 0/8 |
| 27 | H | 307 | + | *** | 8/8 |
| 28 | F | 307 | + | **** | 8/8 |
| 29 | B | 307 | + | * | 4/8 |
| 30 | Td | 307 | + | NS | 1/8 |
| 31 | H | 311 | + | - | 0/8 |
| 32 | F | 311 | + | - | 0/8 |
| 33 | B | 311 | + | - | 0/8 |
| 34 | Td | 311 | + | * | 5/8 |

**Figure 3.**
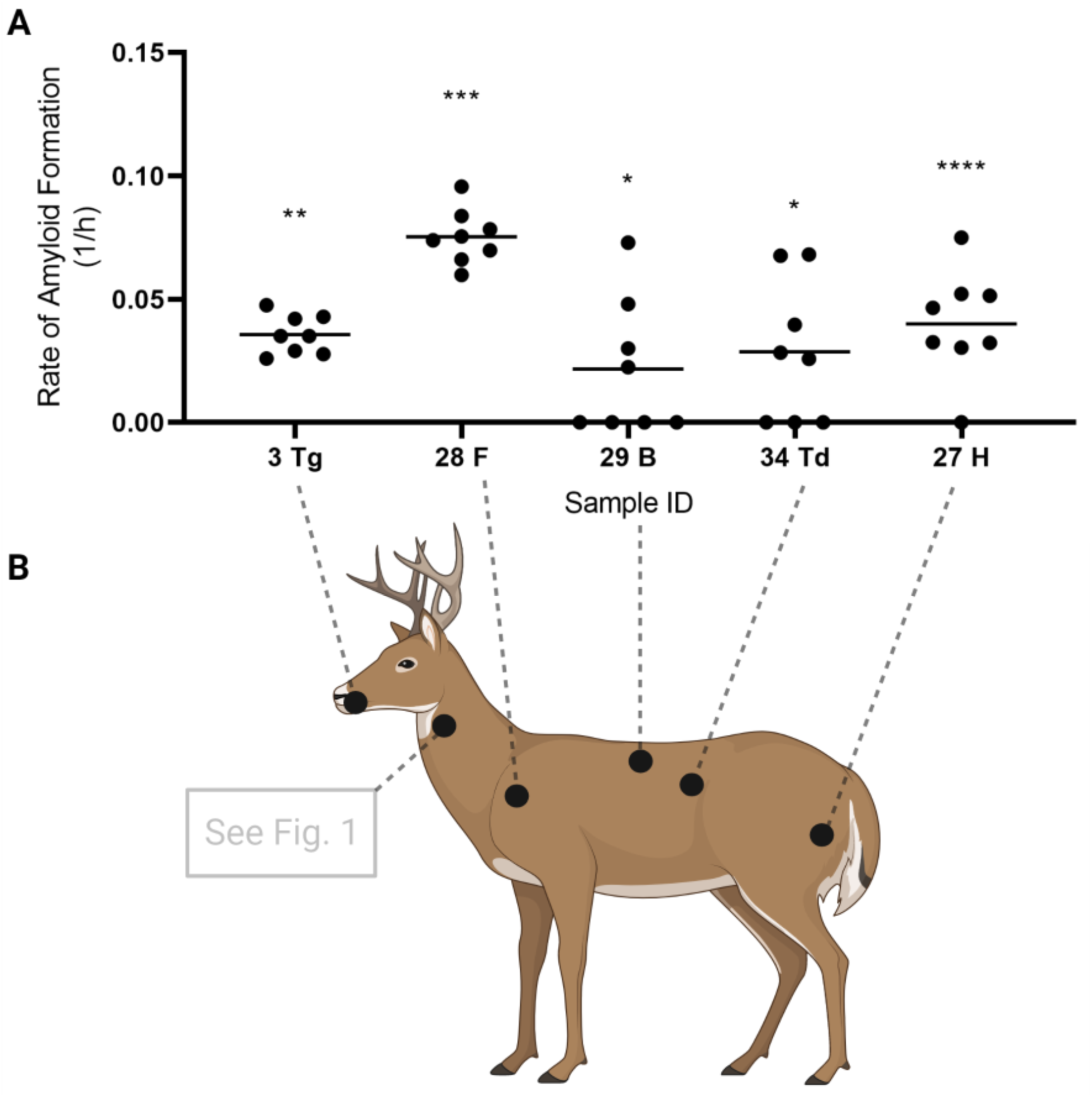
Presence of CWD prions in the muscle tissues of tongue, neck, hindlimb, forelimb, backstrap, and tenderloin. (A) Examples of the rate of amyloid formation (RAF) from RT-QuIC positive samples were plotted. Statistical results as compared to the respective negative controls were indicated (****, p < 0.0001; ***, p < 0.001; **, p < 0.01; *, p < 0.05). (B) Prion-seeding activity was detected by RT-QuIC in muscles from the tongue, forelimb, backstrap, tenderloin, and hindlimb.

Notably, the samples used for this experiment were under various degrees of autolysis. Hypothesizing that this may influence RT-QuIC’s ability to detect prion-seeding activity by changing the optimal dilutions of the processed homogenate, we again looked at prion-seeding activities using serial dilutions of a selected number of samples. As expected, the dilution with adequate positive wells for samples no longer consistently converged at 10^−1^ (Fig 4A), suggesting that the freeze-thaw method is not suitable for muscle tissue samples of sub-optimal quality. To investigate whether other tissue processing methods would improve the detection of CWD prion-seeding activity in RT-QuIC, given the sub-optimal tissue preservation described above, we examined a subset of samples using enzymatic digestions (collagenase A and trypsin) instead of the freeze-thaw method. We hypothesized that collagenase A and/or trypsin would sufficiently digest potential inhibitors to a degree where extensive dilution of the processed homogenates was unnecessary. Surprisingly, collagenase A digestion still required a 10-fold dilution similar to the freeze-thaw method (Supplementary Fig 2) although it appeared to be more sensitive (i.e., identified more muscle samples as RT-QuIC positive from CWD positive animals (Fig 4B)). However, this was not observed when we re-tested a subset of neck muscle samples; in addition, we confirmed that the collagenase method did not appear to produce false positive RT-QuIC signals (Supplementary Fig 3). Alternatively, trypsin digestion produced an extremely high RAF without requiring the 10-fold dilution even though its sensitivity did not improve upon the freeze-thaw method in the given sample set (Fig 4C).

**Figure 4.**
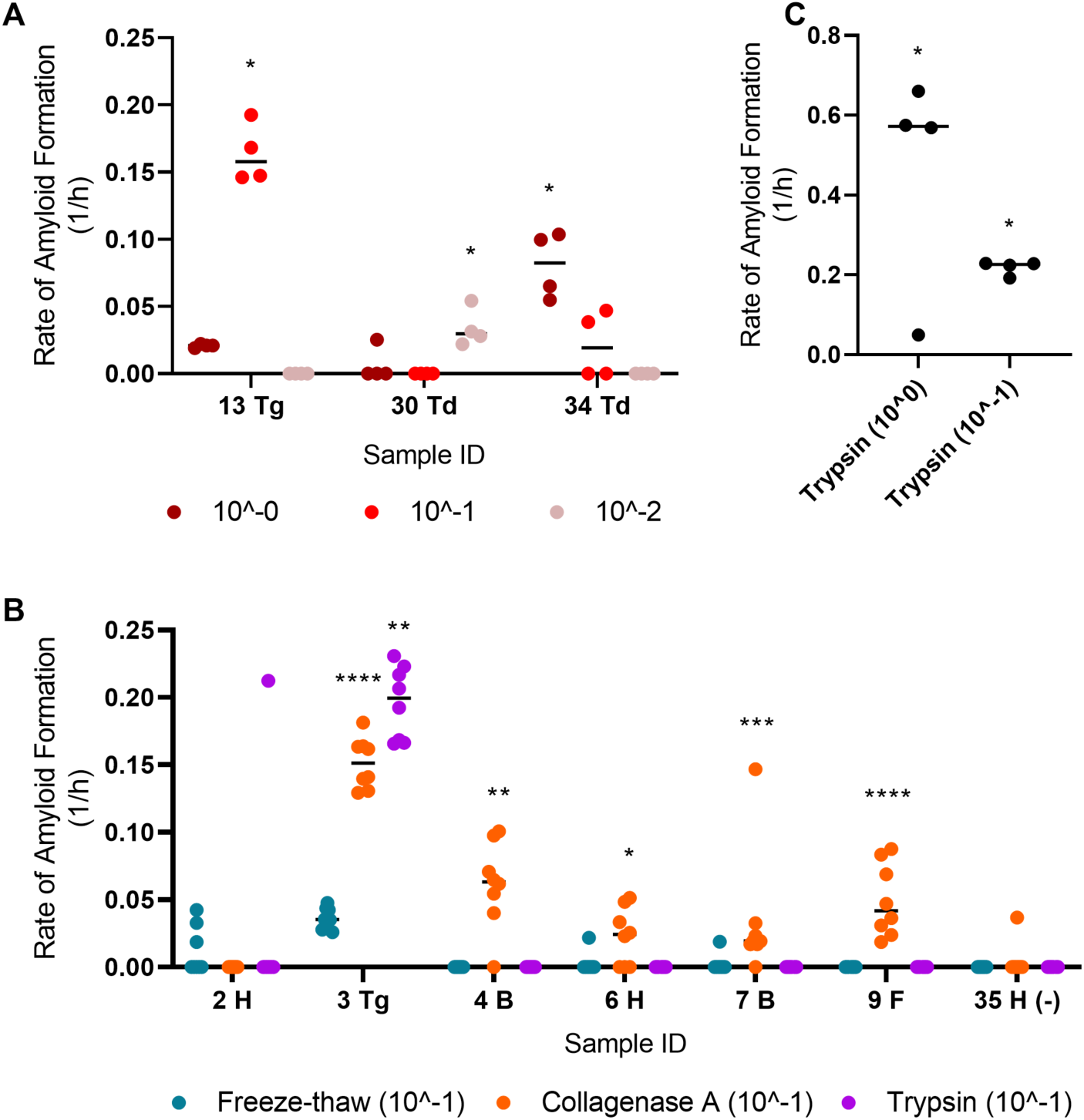
Comparison of different methods used to extract CWD prions from skeletal muscles of WTD. (A) Rate of amyloid formation (RAF) of samples diluted to different concentrations was visualized. (B) RAF from RT-QuIC was plotted for a subset of samples treated by different extraction methods, including freeze-thaw, collagenase A, and trypsin. The final suspension was diluted to 10^-1. (C) RAF of trypsin-digested sample number 3 (tongue; undiluted and diluted to 10^-1) was diagramed. Statistical result for each sample when compared with its respective negative control was indicated (****, p < 0.0001; ***, p < 0.001; **, p < 0.01; *, p < 0.05). B, backstrap; F, forelimb; H, hindlimb; Td, tenderloin; Tg, tongue

We note that all methods used in this study did result in positive prion-seeding activities using RT-QuIC on muscle tissue (Fig 4B). The results reported here indicate that pure mechanical means of extraction may not be enough to facilitate RT-QuIC detection of CWD prions in aged or decomposing muscles.

## Discussion

CWD is an emerging infectious prion disease currently affecting cervid populations across three continents and negatively influencing all cervid-related industries within impacted regions. Infected animals can remain asymptomatic for months while shedding CWD prions through excreta [11,14], thus making the identification of early-stage CWD-infected animals based on external diseased phenotypes impossible. Antibody-based ELISA and IHC tests are the current diagnostic standards for CWD. Despite their reliability, such immunodetection methods have limited sensitivity and application across various tissues and body excreta in comparison to *in vitro* amplification methods for prion detection, such as PMCA and RT-QuIC [5]. Of the available *in vitro* amplification methods, RT-QuIC is ideally suited as a CWD screening tool because it can be easily scaled-up as required by industrial applications. Given the continued spread of CWD, and uncertainty surrounding potential health risks to both animals and humans due to the consumption of CWD-positive venison [1], it is clear that a highly sensitive and reliable diagnostic method to detect CWD prions in skeletal muscles of cervids is needed.

Previous studies indicated that CWD prions were present in the tongue of cervidized transgenic mice and can sustain propagation or infectivity in skeletal muscles from the hindlimb [16,17,27,28]. In this study, we tested methods aimed at extracting and enriching PrP^CWD^ from WTD skeletal muscles for prion detection by RT-QuIC. We first found that CWD prions were present in bulk sampled neck muscles (*brachiocephalicus/sternocephalicus*) of CWD positive animals (Fig 1A) [25]. This result prompted us to investigate the general distribution of prions in skeletal muscles from the tongue, forelimb, mid-truck, and hindlimb of CWD positive WTD tissue-sets available in our biorepository. We found that in addition to the neck, PrP^CWD^ are also present in a variety of skeletal muscle tissues (described above; Table 2; Fig 3). Our results indicate that CWD prions are more widely distributed in skeletal muscles than previously shown.

It remains to be determined whether CWD prions are detectable in skeletal muscles not sampled or in similar studies using amplification-based methods, such as PMCA and RT-QuIC. Although we were unable to detect PrP^CWD^ in every muscle type from CWD positive animals, this result is expected because the successful detection of PrP^CWD^ within an infected individual, and particular tissue type, can be impacted by multiple factors. With respect to the neck samples screened here, we only had access to unilaterally sampled muscles harvested from individual WTD heads and it has been shown recently that PrP^CWD^ may not be bilaterally present in select tissues [25,29]. The 10 positive animals selected for testing were strongly positive across multiple tissues and likely far along in the disease progression [25]. The stability of prions may vary depending on strains [27] and the RT-QuIC method utilized here was not conducted in such a way as to detect particular CWD strain differences. Further, the progression of CWD affects the deposition of prions in peripheral tissues and it is unclear at what time in the disease progression that prions accumulate in WTD muscles. We note the neck muscles were frozen less than 12h after collection however, the other tissue types were at various stages of decomposition and underwent multiple freeze-thaw cycles prior to our having access to them. This difference in tissue preservation and quality likely explains the reduced sensitivity of RT-QuIC upon application, an observation that indicates the accumulation of RT-QuIC inhibitors in decaying tissue.

Based on the results from well-preserved neck muscles, we posit that the freeze-thaw method has the most potential for large-scale diagnostic screening of venison, as it is cheaper and easier to perform. For samples with heavy prion loads, such as tongue, all methods used in this study agreed on the positivity of prion-seeding activity. For poorly preserved sample types, collagenase A outperformed the freeze-thaw method and trypsin digestion in terms of identifying more RT-QuIC positive muscle samples from CWD positive animals. Surprisingly, trypsin digestion yielded a high RAF and did not require additional dilutions of the final resuspension as needed by other methods. This could be due to the digestion of protein inhibitors by trypsin. Additional optimization of the methods presented here is needed for protocols focused on suboptimal sample types. It is possible that the prion seeding activity we detected in the collected muscle tissues is from non-muscle cell types as reported by Daus and colleagues [17]. However, the cellular origin of PrP^CWD^ in skeletal muscle, whether in myocytes, erythrocytes, neurons, epithelial cells, or any other cell type, is inconsequential to the recommendations of not consuming venison from CWD-positive animals or the potential for RT-QuIC-based venison screening as venison products are a matrix of multiple tissues and cell types.

Our study provides the foundation for the development of RT-QuIC-based screening of venison and venison-related products for CWD-prions. Our findings suggest that CWD prions are more widely distributed in the WTD body than previously reported and further investigation is needed to understand the extent of this distribution. Future studies focusing on larger sample sizes with bilateral samplings of well-preserved muscle samples are needed to assess, validate, and improve the presented method for its application, as well as quantify the load of CWD prions present. Longitudinal characterization of prion deposition in a variety of high-quality muscle samples, such as those conducted for saliva, lymphoid tissues, and feces is needed to better understand the pathophysiology of CWD in deer and other cervids. From the perspective of the zoonotic potential of CWD, our research helps reinforce that venison from CWD-positive animals should not be consumed and establish the utility potential of RT-QuIC as a viable method for high-throughput screening of venison products for CWD, thus limiting potentially infectious prions from entering both human and animal food-chains.

## Methods

### Sample collection

RT-QuIC protocol development was initially performed utilizing neck muscle (*brachiocephalicus/sternocephalicus*) tissue samples collected from wild WTD through agency culling operations in southeast Minnesota conducted by the Minnesota Department of Natural Resources in conjunction with USDA APHIS Wildlife Services as described by Schwabenlander et al. [25]. Muscle tissue samples for the quantitative comparison study were obtained through disposal or necropsy of farmed and wild WTD at the University of Minnesota Veterinary Diagnostic Laboratory. All farmed and wild WTD examined in our study have been independently tested for CWD infection based on immunodetection analysis (ELISA and/or IHC) of the brain and/or lymphatic tissue for the presence of PrP^CWD^.

### Muscle preparation

#### Freeze-Thaw method

This method was inspired by a combination of existing RT-QuIC protocols [14,17,30]. Muscles were stored at −20°C within 12h after collection then transferred to −80°C until tested. 10% (weight/volume) muscle homogenates in 1X PBS were prepared in tubes with 1.5 mm diameter zirconium oxide beads using a Beadbug homogenizer at top speed for 180 seconds. The homogenates underwent three cycles of flash freeze-thaw consisting of 3 minutes in dry ice and 3 minutes at 37 °C. The homogenates were subjected to additional homogenization at top speed for 180s using the Beadbug homogenizer. The mixtures were then centrifuged at 5,000 rpm for 3 minutes. 500 μl of supernatants were used for centrifugation at 15,000 rpm, 4°C for 40 minutes. The resultant pellets were resuspended in 100 μl of 1X PBS then incubated with 7 μl of 4% (w/v) phosphotungstic acid (Sigma-Aldrich) in 0.2 M magnesium chloride. The mixtures were then incubated at 37°C and 1500 rpm for 1h in a ThermoMixer (Eppendorf) before being subjected to centrifugation for 30 minutes at 15,000 rpm, 4°C. Pellets were resuspended in 10 μl of 0.1% (v/v) SDS/PBS/N2. 2 μl of 10-1 diluted resuspension was used for optimal result.

#### Collagenase A and trypsin digestion method

This method for RT-QuIC was modified from the PMCA method developed by Daus and colleagues [17].10% (weight/volume, w/v) muscle homogenates and 180-second homogenization were carried out as described above. 350 μl homogenates were mixed with equal volume of 2X collagenase A [4 mM CaCl2 and 0.5% (w/v), Roche] or trypsin (Gibco) solutions. The mixture was incubated at 37°C, 700 rpm for four hours. After being homogenized again for 90 seconds, the mixtures were centrifuged at 5,000 rpm for 3 minutes at 4°C. The supernatant was then transferred to another tube and mixed with an equal volume of 2X protease inhibitor cocktail (Sigma-Aldrich). This was followed by steps including centrifugation at 15,000 rpm for 40 minutes as the freeze-thaw method. 2 μl of the final suspensions were diluted 10-fold and added to the RT-QuIC reaction for Collagenase A digestion. The additional dilution was not necessary for trypsin digestion.

#### RT-QuIC substrate preparation and reaction conditions

Recombinant hamster PrP (HaPrP90-231; provided by NIH Rocky Mountain Laboratory) production and filtration followed the methods of Schwabenlander et al. [25]. An RT-QuIC master mix was made to the following concentrations: 1X PBS, 500 uM EDTA, 50 uM ThT, 300 mM NaCl, and 0.1 mg/mL HaPrPrP. All reagents were filter-sterilized through 0.22um PVDF filters. 98 uL of the master mix was pipetted into each well on a black 96-well plate with clear bottoms. After samples were added, the plate was sealed with clear tape. Plates were shaken on four BMG FLUOstar® Omega microplate readers (BMG LABTECH Inc., Cary, North Carolina, USA) at 700 rpm, double orbital for 57 sec, and then rested for 83 sec. This shake/rest cycle repeated 21 times, then the fluorescence was recorded. Readings were recorded with an excitation filter of 450 nm and an emission filter of 480 nm. The gain was set to 1600. The machine performed 21 flashes/well. No well-scan was conducted. 45°C, 50°C, and 55°C were used. 55°C was only used for investigating whether decomposing tissues would have converging dilutions. 50°C was used for enzymatic digestions.

### Experimental design

The aforementioned RT-QuIC muscle protocols (freeze-thaw and enzymatic digestion) were initially used on a small subset of CWD positive and not-detected neck muscle tissue samples. After refining our methods, we then tested the protocol on a larger set of neck muscles from ten CWD positive and ten CWD not detected deer, with CWD status determined by ELISA, IHC, and RT-QuIC analyses on lymphoid tissues from the animals reported by Schwabenlander et al. [25]. To blind investigators, researcher “A” subsampled approximately 300 mg of each sample, placed them individually in 1.5 ml tubes, and re-labeled them in a randomized numerical order. Researcher “B” carried out the muscle processing and RT-QuIC and was blinded to the original identity of the samples. ELISA and IHC results for animals examined herein are presented in Supplementary Table 1. Researcher “B” was unblinded after the first pass of all samples; investigation of different extraction methods was done after unblinding.

### Data analysis

Statistical analysis and plotting of fluorescence data from RT-QuIC were conducted using GraphPad Prism software (version 9). RT-QuIC data from four or eight replicates were used for calculating the rate of amyloid formation (RAF) for muscles, which is defined by the inverse of the time to reach the fluorescent threshold [10]. The threshold was calculated as ten standard deviations above the average baseline fluorescence unless otherwise specified. We observed variable RAF values across the four microplate readers used for our analyses (i.e. plate reader one consistently exhibited earlier amyloid seeding rates vs. plate readers two, three, and four). In no instance did this impact our positive or negative controls. However, due to RAF differences among plate readers, this threshold could not be applied to all plates. In these rare circumstances, the threshold was calculated as two times the background fluorescence in each well. The differences in RAF calculated by these two methods for a true RT-QuIC positive sample is usually less than 0.01; therefore not influencing general comparisons of RAF among plates. The one-tailed Mann-Whitney unpaired t-test was used to test the average difference between samples and corresponding negative controls on the same plates. Quantitative analysis of CWD prion load in muscles was conducted as described by Henderson and colleagues [10].

## Acknowledgments

We thank Fred Schendel, Tom Douville, and the staff of the University of Minnesota Biotechnology Resource Center for critical support with respect to the large-scale production of recombinant proteins. Evan Kipp and Suzanne Stone provided key assistance with laboratory techniques and logistics. We thank Lon Hebl for graciously providing access to animals housed at the Oxbow Park & Zollman Zoo. We are grateful to the following persons for their assistance with the collection of biological samples used herein: MNDNR Wildlife staff, Roxanne J. Larsen, Negin Goodarzi, Devender Kumar, the Minnesota Board of Animal Health, USDA APHIS Wildlife Services staff, and University of Minnesota Veterinary Diagnostic Lab Necropsy staff, especially Melissa Wolfe. The Minnesota Supercomputing Institute provided secure data storage of computational products stemming from our work. Funding was provided by the Minnesota State Legislature through the Minnesota Legislative-Citizen Commission on Minnesota Resources (LCCMR), Minnesota Agricultural Experiment Station Rapid Agricultural Response Fund, University of Minnesota Office of Vice President for Research, and start-up funds awarded to PAL through the Minnesota Agricultural, Research, Education, Extension and Technology Transfer (AGREETT) program. Fig 3 was organized and partially created using BioRender (biorender.com).

## References

1. Osterholm MT, Anderson CJ, Zabel MD, Scheftel JM, Moore KA, Appleby BS. Chronic Wasting Disease in Cervids: Implications for Prion Transmission to Humans and Other Animal Species. MBio. 2019;27;10(4):e01091–19. PMID: 31337719

2. Prusiner SB. Novel proteinaceous infectious particles cause scrapie. Science. 1982;216(4542):136–44. PMID: 6801762

3. Prusiner SB. Creutzfeldt-Jakob disease and scrapie prions. Alzheimer Dis Assoc Disord. 1989;3(1–2):52–78. PMID: 2568118

4. Ford MJ, Burton LJ, Morris RJ, Hall SM. Selective expression of prion protein in peripheral tissues of the adult mouse. Neuroscience. 2002;113(1):177–92. PMID: 12123696

5. Haley NJ, Richt JA. Evolution of Diagnostic Tests for Chronic Wasting Disease, a Naturally Occurring Prion Disease of Cervids. Pathog. 2017;6(3). PMID: 28783058

6. Kramm C, Soto P, Nichols TA, Morales R. Chronic wasting disease (CWD) prion detection in blood from pre-symptomatic white-tailed deer harboring PRNP polymorphic variants. Sci Rep. 2020;10(1):1–8. PMID: 33188252

7. Weber P, Giese A, Piening N, Mitteregger G, Thomzig A, Beekes M, et al. Cell-free formation of misfolded prion protein with authentic prion infectivity. Proc Natl Acad Sci. 2006;103(43):15818–15823. PMID: 17030802

8. Wilham JM, Orrú CD, Bessen RA, Atarashi R, Sano K, Race B, et al. Rapid End-Point Quantitation of Prion Seeding Activity with Sensitivity Comparable to Bioassays. PLOS Pathog. 2010;6(12):e1001217. PMID: 21152012

9. Atarashi R, Sano K, Satoh K, Nishida N. Real-time quaking-induced conversion: a highly sensitive assay for prion detection. Prion. 2011;5(3):150–3. PMID: 21778820

10. Henderson DM, Davenport KA, Haley NJ, Denkers ND, Mathiason CK, Hoover EA. Quantitative assessment of prion infectivity in tissues and body fluids by real-time quaking-induced conversion. J Gen Virol. 2015;96(1):210–9. PMID: 25304654

11. Henderson DM, Denkers ND, Hoover CE, Garbino N, Mathiason CK, Hoover EA. Longitudinal Detection of Prion Shedding in Saliva and Urine by Chronic Wasting Disease-Infected Deer by Real-Time Quaking-Induced Conversion. J Virol. 2015;15;89(18):9338–9347. PMID: 26136567

12. Hoover CE, Davenport KA, Henderson DM, Denkers ND, Mathiason CK, Soto C, et al. Pathways of Prion Spread during Early Chronic Wasting Disease in Deer. J Virol. 2017;91(10). PMID: 28250130

13. Cooper SK, Hoover CE, Henderson DM, Haley NJ, Mathiason CK, Hoover EA. Detection of CWD in cervids by RT-QuIC assay of third eyelids. PLoS One. 2019;28;14(8):e0221654. PMID: 31461493

14. Tennant JM, Li M, Henderson DM, Tyer ML, Denkers ND, Haley NJ, et al. Shedding and stability of CWD prion seeding activity in cervid feces. PLoS One. 2020;15(3):e0227094. PMID: 32126066

15. Otero A, Duque Velásquez C, Johnson C, Herbst A, Bolea R, Badiola JJ, et al. Prion protein polymorphisms associated with reduced CWD susceptibility limit peripheral PrP(CWD) deposition in orally infected white-tailed deer. BMC Vet Res. 2019;15(1):50. PMID: 30717795

16. Jewell JE, Brown J, Kreeger T, Williams ES. Prion protein in cardiac muscle of elk (*Cervus elaphus nelsoni*) and white-tailed deer (*Odocoileus virginianus*) infected with chronic wasting disease. J Gen Virol. 2006;87(11):3443–50. PMID: 17030881

17. Daus ML, Breyer J, Wagenfuehr K, Wemheuer WM, Thomzig A, Schulz-Schaeffer WJ, et al. Presence and seeding activity of pathological prion protein (PrPTSE) in skeletal muscles of white-tailed deer infected with chronic wasting disease. PLoS One. 2011;6(4):1–7. PMID: 21483771

18. Angers RC, Browning SR, Seward TS, Sigurdson CJ, Miller MW, Hoover EA, et al. Prions in skeletal muscles of deer with chronic wasting disease. Science. 2006;311(5764):1117. PMID: 16439622

19. Barria M, Libori A, Mitchell G, Head M. Susceptibility of Human Prion Protein to Conversion by Chronic Wasting Disease Prions. Emerg Infect Dis J. 2018;24(8):1482. PMID: 30014840

20. Hannaoui S, Schatzl HM, Gilch S. Chronic wasting disease: Emerging prions and their potential risk. PLoS Pathog. 2017;13(11):e1006619. PMID: 29095921

21. FDA. Use of Material from Deer and Elk in Animal Feed. [cited 2021 Feb 6]. Available from: http://www.fda.gov/media/69936

22. FSIS U. Voluntary Inspection of Cervid Animals Tested for Chronic Wasting Disease. 2020.

23. Olszowy KM, Lavelle J, Rachfal K, Hempstead S, Drouin K, Darcy JM, et al. Six-year follow-up of a point-source exposure to CWD contaminated venison in an Upstate New York community: risk behaviours and health outcomes 2005–2011. Public Health. 2014;128(9):860–8. PMID: 25225155

24. Wisconsin Department of Natural Resources. Wisconsin Department of Natural Resources. [cited 2021 Mar 16]. Available from: https://dnr.wisconsin.gov/

25. Schwabenlander MD, Rowden GR, Li M, LaSharr K, Hildebrand EC, Stone S, et al. Comparison of Chronic Wasting Disease Detection Methods and Procedures: Implications for Free-Ranging White-Tailed Deer (*Odocoileus Virginianus*) Surveillance and Management. bioRxiv. 2021;2021.03.03.433751.

26. Bosque PJ, Ryou C, Telling G, Peretz D, Legname G, DeArmond SJ, et al. Prions in skeletal muscle. Proc Natl Acad Sci. 2002;99(6):3812–3817. PMID: 12224617

27. Angers RC, Kang H-E, Napier D, Browning S, Seward T, Mathiason C, et al. Prion Strain Mutation Determined by Prion Protein Conformational Compatibility and Primary Structure. Science. 2010;328(5982):1154–1158. PMID: 20466881

28. Seelig DM, Mason GL, Telling GC, Hoover EA. Pathogenesis of Chronic Wasting Disease in Cervidized Transgenic Mice. Am J Pathol. 2010;176(6):2785–97. PMID: 20395435

29. Bloodgood J, Kiupel M, Melotti J, Straka K. Chronic Wasting Disease Diagnostic Discrepancies: The Importance of Testing Both Medial Retropharyngeal Lymph Nodes. J Wildl Dis. 2021;57(1):194–8. PMID: 33635974

30. Elder AM, Henderson DM, Nalls A V, Hoover EA, Kincaid AE, Bartz JC, et al. Immediate and Ongoing Detection of Prions in the Blood of Hamsters and Deer following Oral, Nasal, or Blood Inoculations. J Virol. 2015;89(14):7421–7424. PMID: 25926635

