## Supplemental Information for "RT-QuIC detection of CWD prion seeding activity in white-tailed deer muscle tissues"

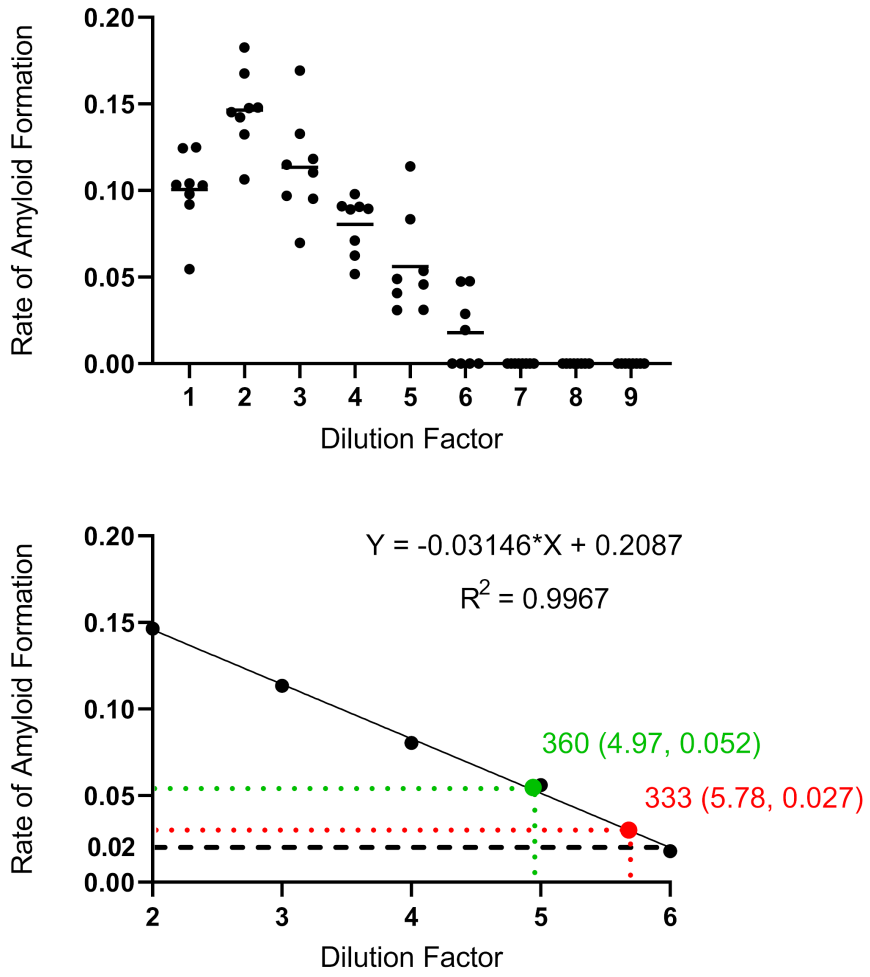


**Supplementary Figure 1. Comparison of RAF (rate of amyloid formation) between lymphoid and skeletal muscle tissues.**

(A) RAF of 10% lymphoid tissue homogenates was plotted against dilution factors ranging from 1 to 9. The RT-QuIC experiment was set at 45°C. 8 replicates were conducted for all dilutions.
(B) The average of RAF from dilutions in A) with dilution factors ranging from 2-6 was fitted on a linear model. Equation and R^2 values are indicated. RAF for neck muscle samples from animal 333 and 360 processed using the freeze-thaw method and normalized to the lymphoid tissue used in A) were plotted.

Result: Prion load using muscles processed by the freeze-thaw method based on RAF from RT-QuIC results is equivalent to 10^-6~10^-7$^{th}$ (2/1000 * 10/2 * 10 * 10^-5~10^-6) that of lymphoid tissues.

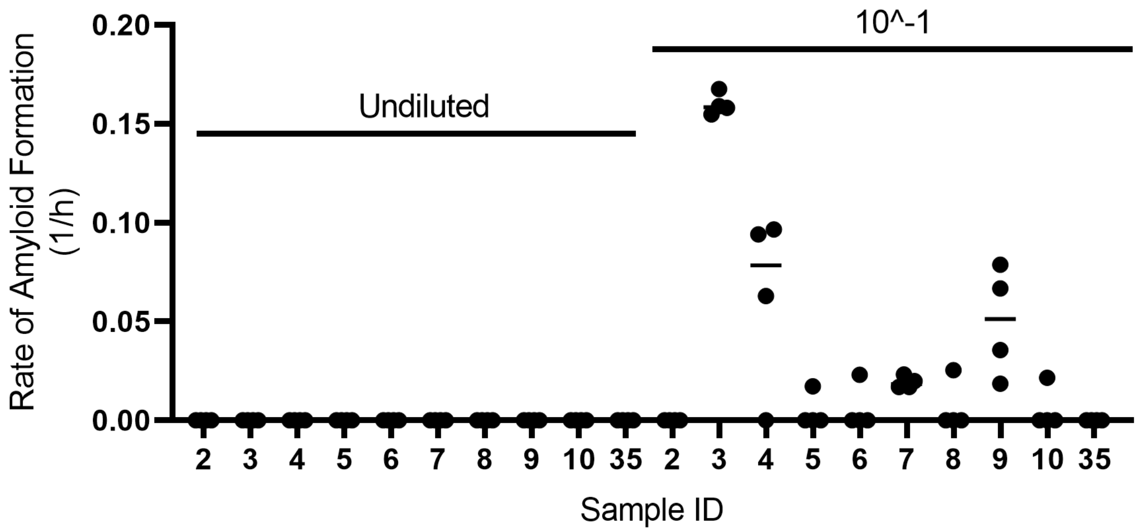


**Supplementary Figure 2. Collagenase A-processed samples need to be diluted to facilitate prion-seeding activity detection in RT-QuIC**.
A subset of samples with undiluted final suspension and that diluted to 10^-1 were tested on the same plate.

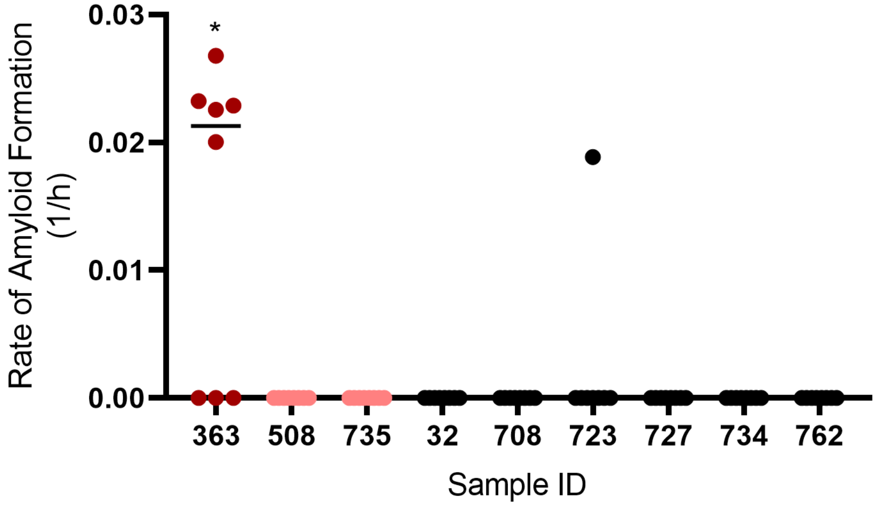


**Supplementary Figure 3. Collagenase A processing does not produce significant RT-QuIC false positives for neck muscles**.
