## Supplementary Table 1 for "RT-QuIC detection of CWD prion seeding activity in white-tailed deer muscle tissues"

| MNPRO ID | Age | Sex | Background | Official CWD Test Results | Tissues Sampled** |
|---|---|---|---|---|---|
| 166 | Adult | Female | Wild | Positive | N |
| 250 | Adult | Male | Wild | Positive | N |
| 333 | Adult | Female | Wild | Positive | N |
| 353 | Adult | Female | Wild | Positive | N |
| 360 | Adult | Female | Wild | Positive | N |
| 363 | Adult | Female | Wild | Positive | N |
| 376 | Adult | Female | Wild | Positive | N |
| 384 | Adult | Female | Wild | Positive | N |
| 508 | Adult | Male | Wild | Positive | N |
| 735 | Adult | Female | Wild | Positive | N |
| 118 | Fawn | Male | Wild | Not detected | N |
| 125 | Yearling | Female | Wild | Not detected | N |
| 239 | Fawn | Male | Wild | Not detected | N |
| 443 | Adult | Female | Wild | Not detected | N |
| 515 | Adult | Female | Wild | Not detected | N |
| 536 | Yearling | Female | Wild | Not detected | N |
| 693 | Yearling | Male | Wild | Not detected | N |
| 708 | Yearling | Female | Wild | Not detected | N |
| 723 | Adult | Female | Wild | Not detected | N |
| 727 | Adult | Female | Wild | Not detected | N |
| 734 | Fawn | Female | Wild | Not detected | N |
| 762 | Adult | Female | Wild | Not detected | N |
| 287 | Adult | Male | Wild | Positive | B, F, H, Tg |
| 288 | Adult | Female | Wild | Positive | B, F, H |
| 289 | Adult | Male | Wild | Positive | B, F, H |
| 290* | Adult | Female | Wild | Positive | B, F, H |
| 295 | Adult | Female | Wild | Positive | F, H, Td, Tg |
| 296 | Adult | Female | Wild | Positive | B, F, H |
| 297 | Yearling | Male | Wild | Positive | B, F, H |
| 298 | Adult | Female | Wild | Positive | B, F, H |
| 307* | Adult | Female | Farmed | Positive | B, F, H, Td |
| 311 | Adult | unknown | Wild | Positive | B, F, H, Td |
